## Supplementary figures and images for "Fast and accurate imputation of genotypes from noisy low-coverage sequencing data in bi-parental populations"

### S1 Fig

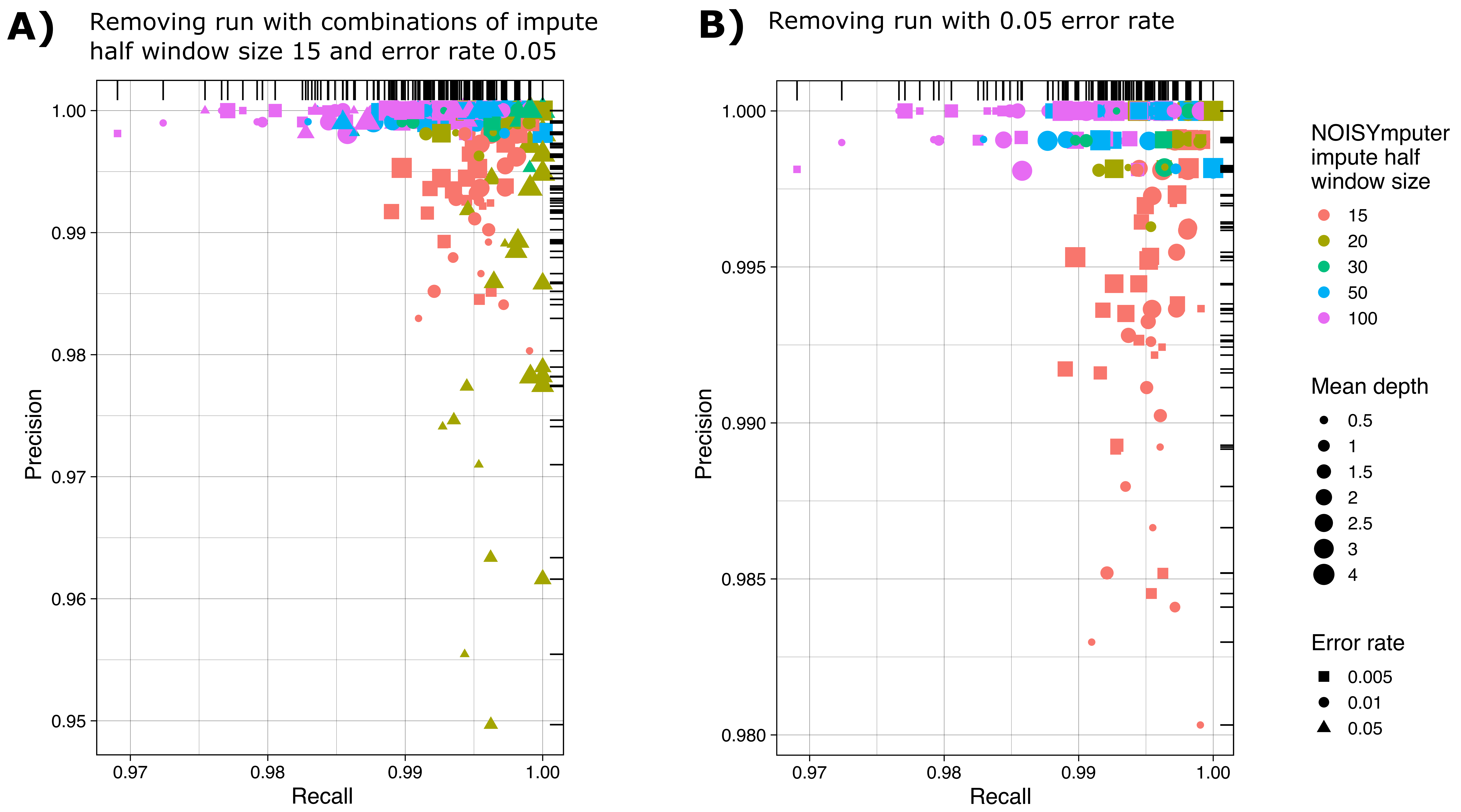
