## Supplementary material for "Fast and accurate imputation of genotypes from noisy low-coverage sequencing data in bi-parental populations": S1 file

### Options used for imputation with LB-impute, TASSEL-FSFHap and NOISYmputer.

**LB-impute**
Options used for Maize and Rice datasets

| ## Parents imputation | ## Offspring imputation |
|---|---|
| -window 5<br>-minsamples 5<br>-minfraction 0.5<br>-recombdist 10000000<br>-genotypeerr 0.05<br>-readerr 0.05<br>-dr<br>-parentimpute | -window 5<br>-minsamples 5<br>-minfraction 0.5<br>-recombdist 10000000<br>-genotypeerr 0.05<br>-readerr 0.05<br>-dr<br>-offspringimpute |

**FSFHap**
Options used for Maize, Rice and simulated datasets
Pedigree file have been changes depending on tested VCFs.

```
-FSFHapImputationPlugin
-pedigrees pedigreefile
-windowLD true
-bc false
-multbc false
-phet 0.5
-merge false
-outParents true
-endPLugin
-exportType VCF
```

**NOISYmputer**
Options used for Maize, Rice and simulated datasets

| ## Maize and Rice datasets | ## Simulated datasets |
|---|---|
| All defaults options except for:<br><br>imputeHalfWindowSize=30<br>errorA=0.005<br>errorB=0.005<br>dropBreakpointSupportInterval=2<br>exportFormat=VCF | All defaults options except for:<br><br>errorA=0.05<br>errorB=0.05<br>alphaFillGaps=0.001<br>exportFormat=VCF<br><br>with imputeHalfWindowSize set to either 15, 20, 30, 50 or 100. |

### Generation of Rice F2 VCF dataset

The $F_2$ rice dataset was produced using a Snakemake pipeline processing data from fastq to VCF along with a conda environment available at:
https://forge.ird.fr/diade/recombination_landscape/biparental-lc-ngs-snakemake/-/tree/main

### Cluster description

We utilized the IFB Core Cluster provided by the Institut Français de Bioinformatique. This infrastructure is a coordinated network of the national IFB-core Cluster and regional platform infrastructures. Jobs were executed on nodes selected by the cluster scheduler based on the requested and available resources. Consequently, all jobs were run on nodes belonging to one of these machines:

- 16 compute nodes (128 cores, 2 TB RAM) HPE Apollo 2000 XL225n Gen10+, 2x AMD EPYC 7662 (3.1GHz, 64 cores), 3 TB RAM

- 68 compute nodes (28 cores, 256 GB RAM) DELL C6320, 2x Intel Xeon E5-2695v3 (2.3GHz, 14 cores), 256GB RAM
